# Apiaceae *FNS I* originated from *F3H* through tandem gene duplication

**DOI:** 10.1101/2022.02.16.480750

**Authors:** Boas Pucker, Massimo Iorizzo

## Abstract

**Background:** Flavonoids are specialized metabolites with numerous biological functions in stress response and reproduction of plants. Flavones are one subgroup that is produced by the flavone synthase (FNS). Two distinct enzyme families evolved that can catalyze the biosynthesis of flavones. While the membrane-bound FNS II is widely distributed in seed plants, one lineage of soluble FNS I appeared to be unique to Apiaceae species.

**Results:** We show through phylogenetic and comparative genomic analyses that Apiaceae *FNS I* evolved through tandem gene duplication of flavanone 3-hydroxylase (*F3H)* followed by neofunctionalization. Currently available datasets suggest that this event happened within the Apiaceae in a common ancestor of *Daucus carota* and *Apium graveolens*. The results also support previous findings that *FNS I* in the Apiaceae evolved independent of *FNS I* in other plant species.

**Conclusion:** We validated a long standing hypothesis about the evolution of Apiaceae FNS I and predicted the phylogenetic position of this event. Our results explain how an Apiaceae-specific *FNS I* lineage evolved and confirm independence from other *FNS I* lineages reported in non-Apiaceae species.

## Introduction

A plethora of specialized metabolites including flavonoids are produced by plants. These compounds provide an evolutionary advantage under certain environmental conditions. Flavonoids are produced in response to stresses like ultra violet (UV) radiation, cold, or drought [1, 2]. Especially visible is the pigmentation of flowers and fruits by anthocyanins which are one subclass of the flavonoids [3, 4]. Other subclasses include the proanthocyanidins which contribute to the pigmentation of seed coats [5] or flavonols which are produced in response to UV stress [6]. These branches of the flavonoid biosynthesis are well studied and conserved in many plant species and represent a model system for the investigation of the specialized metabolism in plants. A less conserved branch of the flavonoid biosynthesis leads to flavones (Fig 1), which are important in signaling and defense against pathogens [7]. Flavones are derivatives of phenylalanine which is channeled through the general phenylpropanoid pathway to the chalcone synthase (CHS). This enzyme is the first committed step of the flavonoid biosynthesis. Chalcone isomerase (CHI) and flavone synthase (FNS) represent the following steps involved in the formation of the flavone apigenin. F3’H can convert naringenin into eriodictyol which serves as substrate for the formation of the flavone luteolin (Fig 1). FNS activity evolved independently in different phylogenetic lineages [8], and to date two types of FNS genes have been described, named FNSI and FNSII. Distributed over a wide phylogenetic range is FNS II, a membrane-bound cytochrome P450-dependent monooxygenase [9]. An independent lineage of FNS I, a soluble Fe^2+^/2-oxoglutarate-dependent dioxygenase (2-ODD), was identified in the Apiaceae and appeared to be restricted to that family [10]. However, other studies report FNS I functionality in other plant species like OsFNSI in *Oryza sativa* [11], EaFNSI in *Equisetum arvense* [12], PaFNSI in *Plagiochasma appendiculatum* [13], AtDMR6 in *Arabidopsis thaliana* [14], and ZmFNSI-1 in *Zea mays* [14]. These lineages were presented as independent evolutionary events and are not orthologs of Apiaceae *FNS I* [8, 15]. Recently, a study revealed that FNS I is widely distributed in liverworts and is the most likely origin of seed plant flavanone 3-hydroxylase (F3H) [8]. Reports of enzymes with multiple functions like F3H/FNS I [8, 10] or F3H/FLS [16, 17] indicate that the 2-ODD family has a high potential for the acquisition of new functionalities and that independent evolution of these new functions might happen frequently.

**Fig 1.**
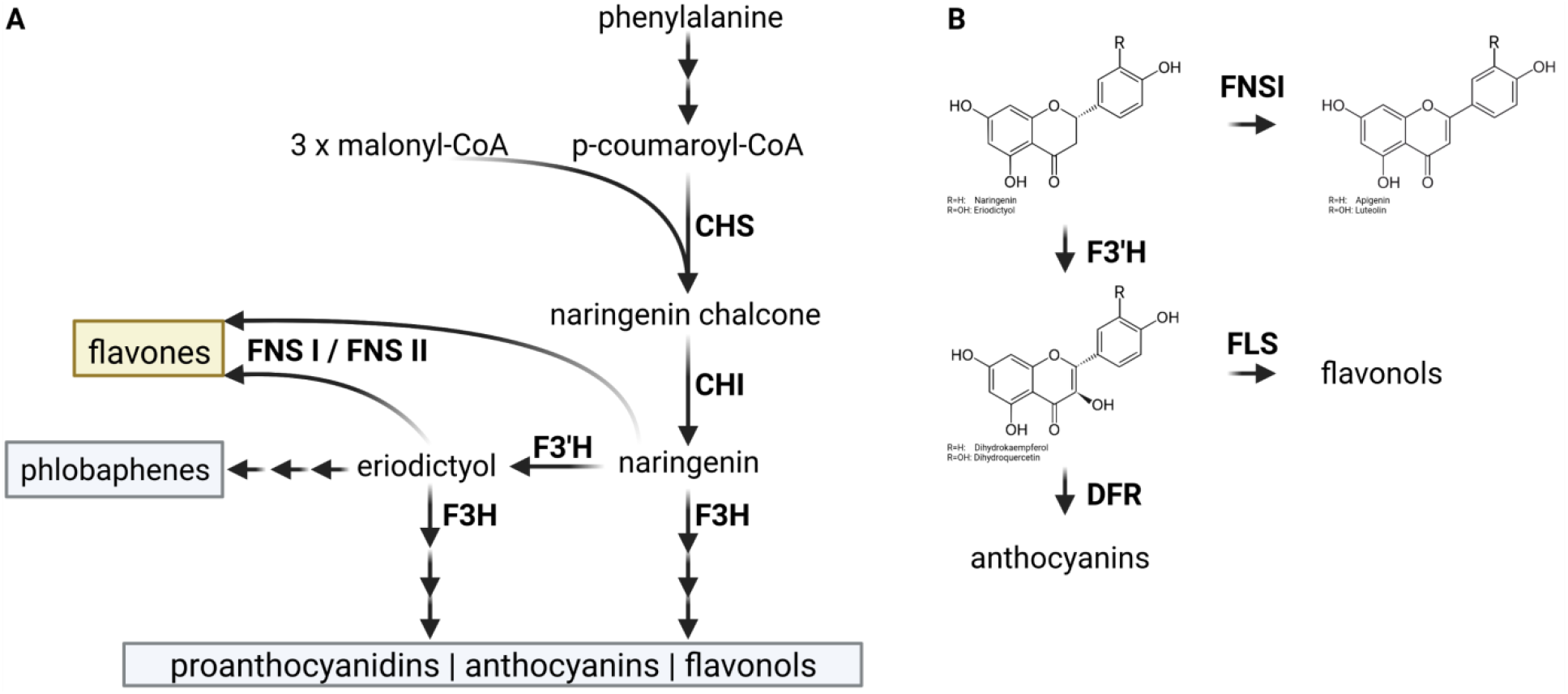
(A) Simplified illustration of the flavonoid biosynthesis with focus on the flavone biosynthesis. CHS (naringenin-chalcone synthase), CHI (chalcone isomerase), F3H (flavanone 3-hydroxylase), F3’H (flavonoid 3’-hydroxylase), and FNS (flavone synthase). (B) Reactions catalyzed by FNS I and F3H in the context of the flavonoid biosynthesis.

It is assumed that flavone biosynthesis is an evolutionarily old trait that predates flavonol and anthocyanin biosynthesis, because the ancestor of the F3H was probably a FNS I [8]. Minor changes in sequence and protein structure can determine the change in enzyme function. One particularly important residue is Y240 in the liverwort *Plagiochasma appendiculatum* PaFNSI/F2H [13]. Bifunctional *Physcomitrella patens* and *Selaginella moellendorffii* enzymes show M or F residues at this site. Most angiosperms and gymnosperms show a P at the corresponding position of their F3Hs [8]. Substitution of this P by M or F resulted in reduced F3H activity and increased FNS I activity, while a replacement with Y resulted in dominant FNS I activity [8]. This indicates that this site played a crucial role in the transition from FNS I to F3H activity.

Apiaceae FNS I show high sequence similarity to F3H thus both were previously classified as DOXC28 in a systematic investigation of the 2-ODD family [18]. Another study called this group of 2-ODD sequences ‘POR’, because they are NADPH-cytochrome P450 oxidoreductases [19]. It was also hypothesized that Apiaceae *FNS I* evolved from *F3H* of seed plants by duplication and subsequent divergence [10, 19, 20]. F3H and FNS I accept the same substrate (Fig 1) which suggests that competition takes place if both enzymes are present at the same intracellular location. The specific activity of both enzymes in the Apiaceae is defined by a small number of diagnostic amino acid residues [8, 10]. It is important to note that the P/Y substitution [8] described above does not play a role in the Apiaceae, because FNS I and F3H sequences show a conserved P at this position. Substitution of several other amino acids in F3H results in FNS I activity though [10]. For instance, I131F, M106T, and D195E are sufficient to confer partial FNS I function to F3H [10]. Also, I131F with L215V and K216R can be sufficient to confer FNS I functionality [10]. A substitution of these seven amino acid residues substantially modifies the pocket of the active site hence changing the orientation of the substrate [10]. This is expected to cause a syn-elimination of hydrogen from carbon-2 (FNS activity) instead of hydroxylation of carbon-3 (F3H activity) [10].

Although previous work hypothesized that Apiaceae FNS I originated from F3H through duplication and neofunctionlization [10, 19, 20], this hypothesis has not yet been validated. The recent release of high quality genome sequences representing most angiosperm lineages including members of the Apiacea family [21–24] opens the opportunity to address this hypothesis. Here, we investigated the evolution of FNS I in the Apiaceae through phylogenetic analysis and comparative genomics. The results indicate that *FNS I* originated from a tandem duplication of *F3H* that was followed by a neofunctionalization event.

## Methods

### Datasets

The genome sequences and the corresponding annotation of *Daucus carota* 388_v2.0 [21] and *Panax ginseng* GCA_020205605.1 [22] were retrieved from Phytozome [25]. The genome sequences of *Apium graveolens* GCA_009905375.1 [23] and *Centella asiatica* GCA_014636745.1 [24] were downloaded from NCBI. Sequences of F3H and FNS I were retrieved from the KIPEs v1 data set [26] and are included in S1 File. The phylogenetic relationships of Apiaceae species were inferred from a previously constructed species tree [27]. This tree was used to arrange genome sequences in the synteny analysis.

### Gene prediction

Since no complete annotation of the coding sequences was publicly available for *Apium graveolens* and *Centella asiatica*, we applied AUGUSTUS v3.3 [28] for an *ab initio* gene prediction with previously described settings [29]. The *Daucus carota* annotation of *F3H* and *FNS I* was manually checked in the Integrated Genomics Viewer [30] and revised (S1 File). Polishing of the gene models was based on a TBLASTN v2.8.1 [31] alignment of the *Petroselinum crispum* FNS I sequence against the *D. carota* genome sequence. Additionally, RNA-seq reads were retrieved from the Sequence Read Archive (S2 File) and aligned to the *D. carota* genome sequence using STAR v2.7.3a [32] with previously described parameters [33].

### Alignment and phylogenetic tree construction

F3H and FNS I polypeptide sequence collections [26] were used to search for additional candidates in *C. asiatica*, *D. carota*, and *A. graveolens* using a BLASTp-based Python script [34]. Initial candidates were validated through a phylogeny constructed with FastTree v2.1.10 (-wag-nosupport) [35] based on a MAFFT v7.475 [36] alignment of the polypeptide sequences. Additional phylogenies were constructed based on the collected polypeptide sequences with different approaches to validate important relationships. MAFFT v7.475 [36] and MUSCLE v5.1 [37] were applied for the alignment construction with default parameters. A customized Python script (algntrim.py, [34]) was applied to remove alignment columns with less than 10% occupancy. The maximum likelihood tree displayed in Fig 2 is based on the MAFFT alignment running RAxML-NG v1.0.1 [38] with LG+G8+F and 10300 rounds of bootstrapping. RAxML-NG was also run with the same model based on a MUSCLE5 alignment of the same polypeptide sequences. Phylogenies for both alignments were also generated with FastTree2 [35] (-wag), IQ-TREE v1.6.12 (-alrt 1000 -bb 1000)[39, 40], and MEGA v11.0.13 [41] (neighbor-joining, 1000 bootstrap replicates, poisson model, uniform rates, pairwise deletion). The resulting tree topologies were manually compared to validate (1) important nodes, (2) monophyly of FNS I, and (3) position of the FNS I clade within the F3H clade of the Apiaceae.

**Fig 2.**
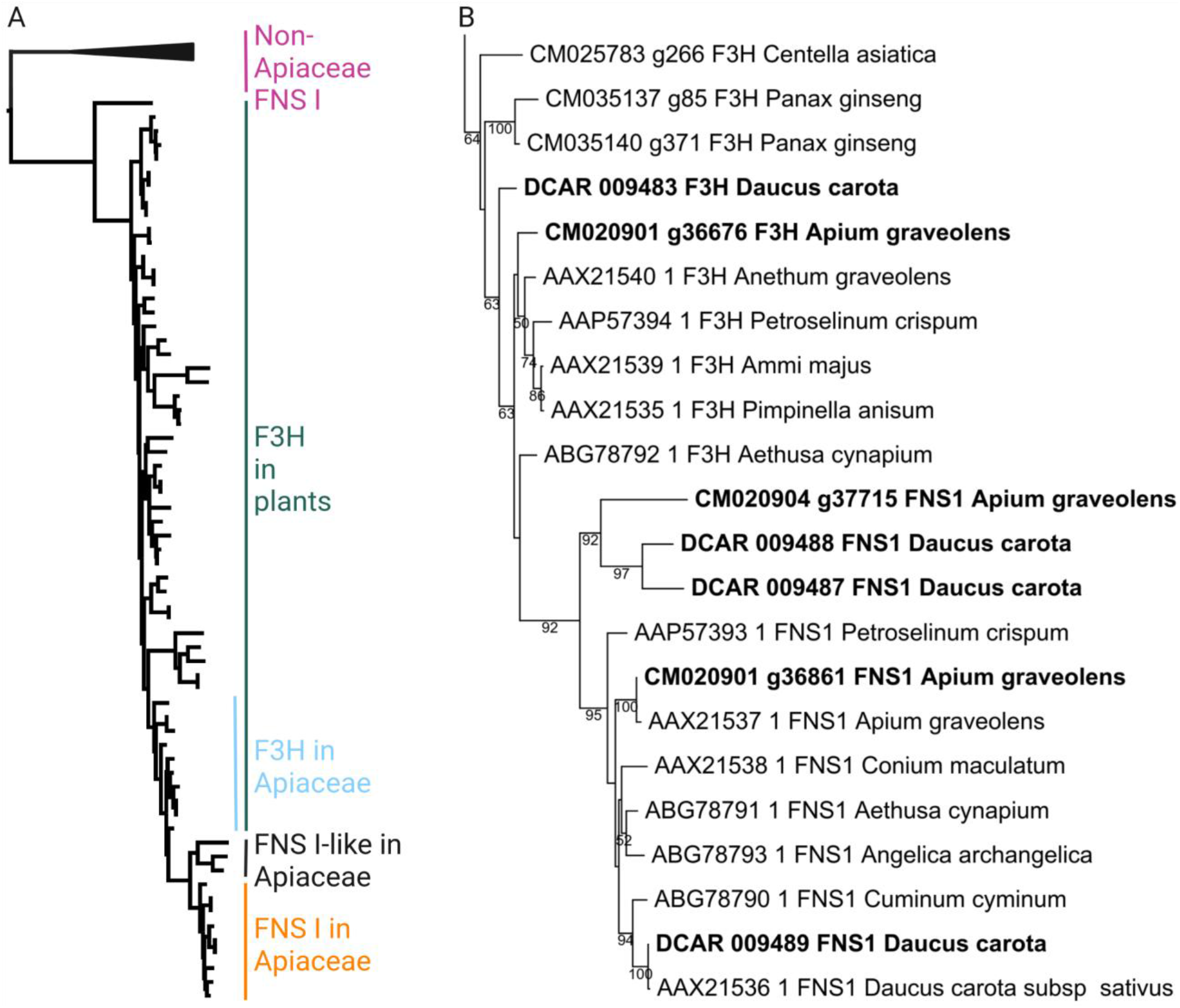
(A) Maximum likelihood tree of F3H and FNS I sequences. Apiaceae FNS I sequences form a nested cluster within the F3H context. The FNS I sequences of non-Apiaceae species are placed outside the F3H clade. Polypeptide sequences used for the construction of this tree are included in S1 File. A full phylogenetic tree with sequence names is included as S3 File. (B) Apiaceae clade in the maximum likelihood tree of F3H and FNS I sequences. Sequences analyzed in Table 1 are written in bold. Classification of sequences in the Apiaceae F3H, Apiaceae FNS I, and Apiaceae FNS I-clades were supported by RAxML, FastTree, MEGA, and IQ-TREE analyses. Support values above 50 are given for the individual branches.

### Synteny analysis

JCVI/MCscan [42] was applied to compare the genome sequences of *P. ginseng*, *C. asiatica*, *D. carota*, and *A. graveolens*. The region around *F3H* and *FNS I* was manually selected. Connections of genes between the species were manually validated and revised based on phylogenetic trees (S3 File). TBLASTN v2.8.1 (-evalue 0.00001) [43] was run with the *P. crispum* FNS I against the genome sequence of *C. asiatica* and *A. graveolens* to identify gene copies that might be missing in the annotation. The results of this search were compared against the annotation to find BLAST hits outside of gene models [34]. The best hits were assessed in a phylogenetic tree with previously characterized F3H and FNS I sequences.

### Gene expression analysis

Paired-end RNA-seq data sets were retrieved from the Sequence Read Archive via fastq-dump v2.8.1 [44] (S2 File). kallisto v0.44 [45] was applied with default parameters for the quantification of gene expression (counts and TPMs). A Python script was developed for the generation of violin plots to illustrate the variation of gene expression (TPMs) across various samples [34]. Outliers, defined as data points which are more than three interquartile ranges away from the median, were excluded from this visualization. Co-expression was analyzed by calculating pairwise Spearman correlation coefficients of gene expression values across all samples. Lowly expressed genes were excluded and only pairs with a correlation coefficient >0.65 and an adjusted p-value < 0.05 were reported. Functional annotation of the *Daucus carota* genes (Dcarota_388_v2.0) was inferred from *Arabidopsis thaliana* based on reciprocal best BLAST hits of the representative peptide sequences as previously described [29, 46]. This co-expression analysis was implemented in a Python script (coexp3.py) which is available via github [34] and as an online service [47].

## Results

Apiaceae FNS I sequences show high similarity to F3H that suggest a close phylogenetic relationship of both lineages. For example, *D. carota* FNS I and F3H have 78% identical amino acids, while the proportion of identical amino acids between different FNS I candidates is 80% (S4 File). A phylogenetic tree was constructed to visualize the relationship of all these sequences in a larger context. FNS I sequences of the non-Apiaceae species *Arabidopsis thaliana*, *Oryza sativa*, *Zea mays*, and *Parmotrema appendiculatum* clustered outside the F3H clade in this tree. The FNS I sequences of seven Apiaceae species formed a distinct clade (Fig 2). This FNS I clade is embedded within a large clade of F3H sequences that included a wide range of phylogenetically distant plants. The position of the FNS I sequences within the F3H clade suggests that Apiaceae *FNS I* originated from *F3H*. The pattern also supports a single *FNS I* origin within the Apiaceae. The critical node separating Apiaceae FNS I from Apiaceae F3H is well supported in the results of all applied tools (S3 File). The monophyly of the Apiaceae FNS I clade is also well supported in all analyses. The FNS I sequences of non-Apiaceae species seem to have an independent origin.

A previous study identified diagnostic amino acid residues that determine the FNS or F3H activity, respectively [10]. It was demonstrated that a substitution of selected amino acid residues can convert one enzyme into the other. We inspected these characteristic features of the FNS I and F3H sequences of *Daucus carota* and *Apium graveolens* (Table 1). The results suggest that there is one *bona fide* F3H in *D. carota* (DCAR_009483) and *A. graveolens* (CM020901_g36676), respectively (S1 File). We also identified one FNS I in each of these species: DCAR_009489 and CM020901_g36861, respectively. In addition, there are FNS I-like copies which lack some of the functionally important amino acid residues of a *bona fide* FNS I (Table 1). The separation of the *FNS I*-like lineage from the *FNS I* lineage seems to predate a duplication in the *FNS I*-like lineage that produced the two copies discovered in *D. carota* and *A. graveolens*.

**Table 1.**
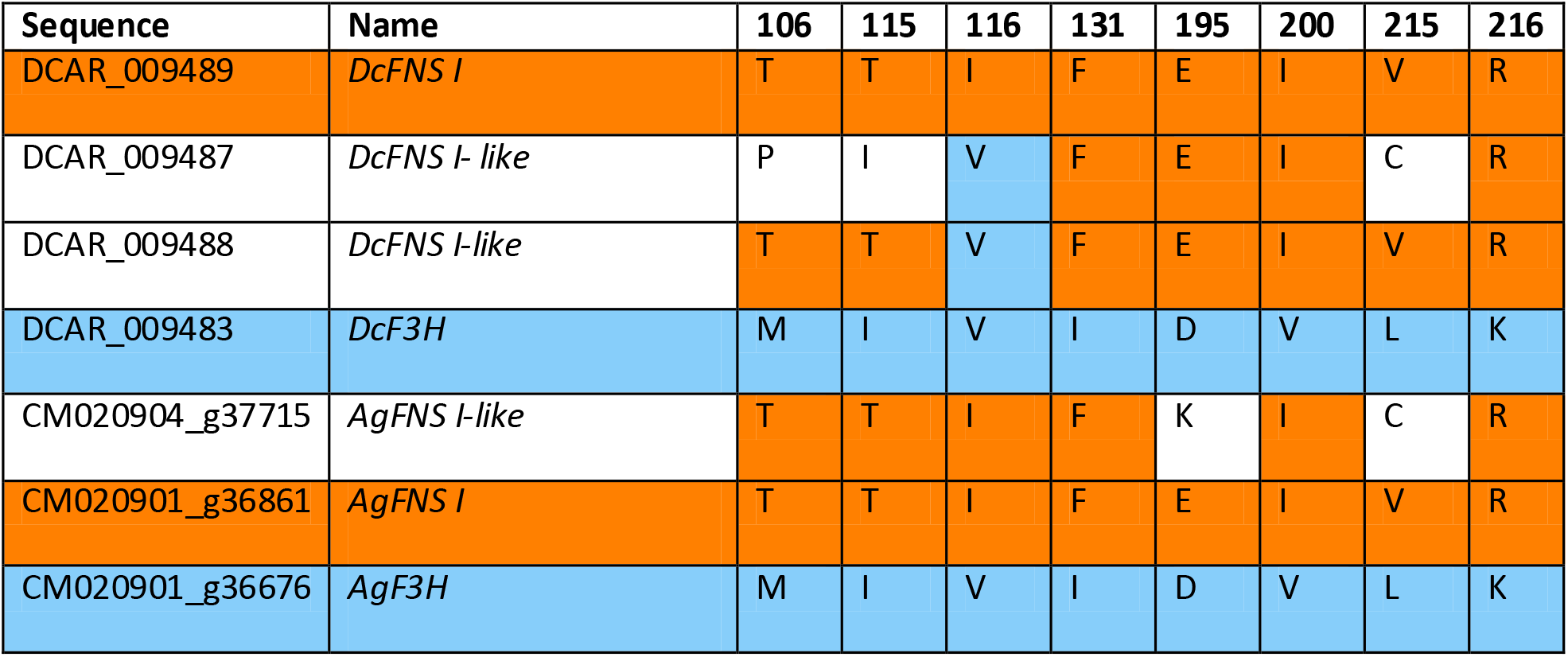
Inspection of diagnostic amino acid residues in FNS I and F3H candidates of *Daucus carota* and *Apium graveolens*. FNS I residues are highlighted in orange, F3H residues are highlighted in skyblue. Positions are based on the FNS I of *Petroselium crispum* (AAP57393.1).

To narrow down the origin of the Apiaceae *FNS I*, we compared highly contiguous genome sequences of Apiaceae and outgroup species. The Apiaceae members *Daucus carota* and *Apium graveolens* show microsynteny in a region that harbors both, *F3H* and *FNS I* genes (Fig 3). Both species differ from the *Centella asiatica* (basal Apiaceae species) and *Panax ginseng* (outgroup species) which do not show a *FNS I* gene in this region or elsewhere in the genome sequence. However, the presence of *F3H* and synteny of many flanking genes indicates that the correct region was analyzed.

**Fig 3.**
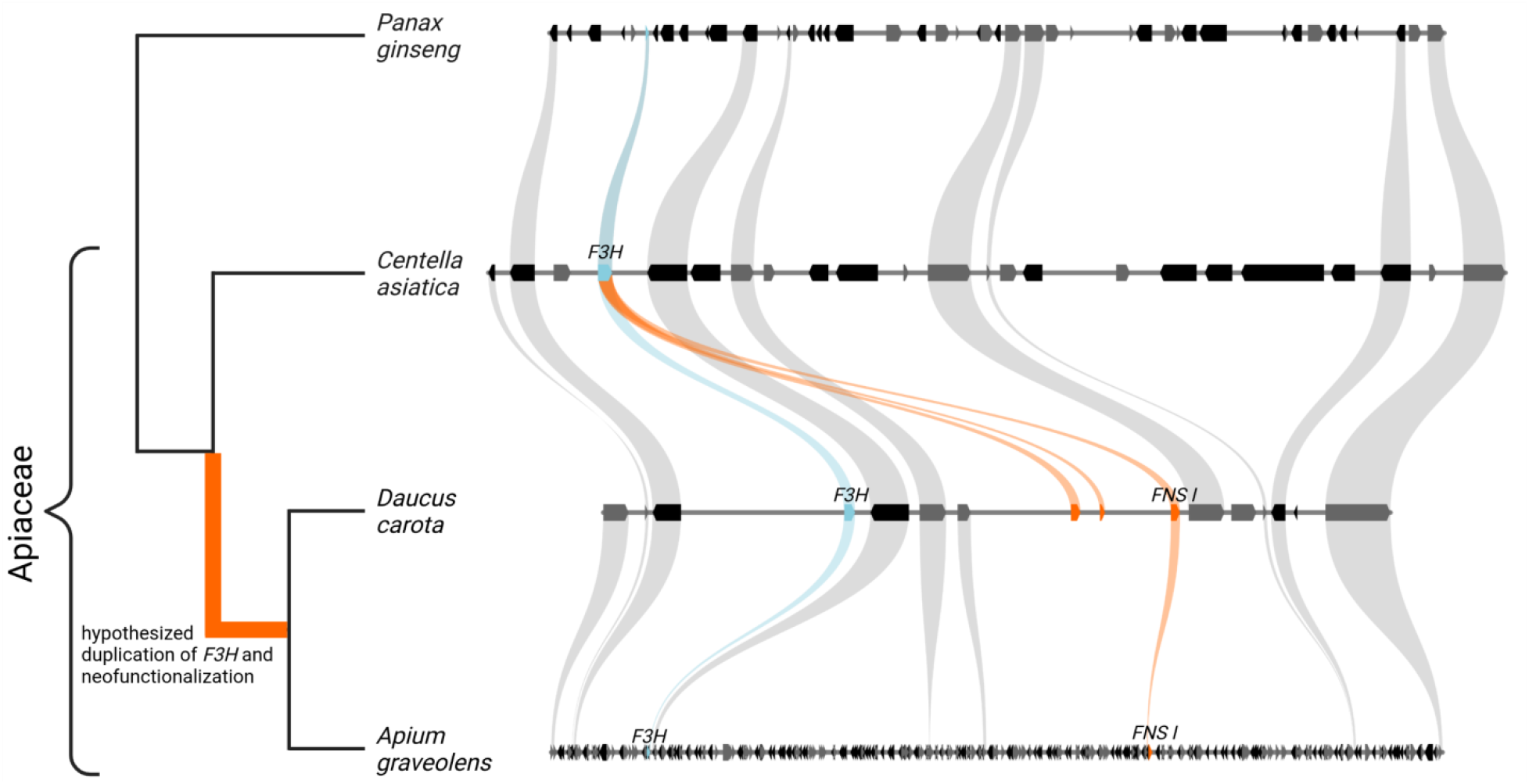
Syntenic region between the Apiaceae species *Daucus carota* and *Apium graveolens* shows *F3H* (skyblue) and *FNS I* (orange) in close proximity, while *FNS I* was not observed in the basal Apiaceae species *Centella asiatica* or in the outgroup *Panax ginseng*.

Although multiple gene copies were identified based on the available genome sequences, expression of these genes determines their relevance. Expression of the *F3H*, *FNS I*, and *FNS I*-like genes in carrots was analyzed across 146 RNA-seq samples (Fig 4). The results show that *F3H* and *FNS I* show substantially higher expression than any of the *FNS I*-like genes (DCAR_009487) while the other *FNS I*-like gene (DCAR_009488) is almost not expressed. Expression analysis in specific plant parts and tissues revealed that *F3H* (DCAR_009483) is strongly expressed in phloem and flowers, while *FNS I* (DCAR_009489) was dominant in leaf, petiole, and root (Fig 4). Expression patterns of both *FNS I*-like genes are more similar to *FNS I* than to *F3H* expression. Strongest expression of DCAR_009487 was observed in the phloem and xylem of the root and in the petiole. DCAR_009488 showed the highest expression in whole flowers and stressed leaves of orange cultivars (S2 File). Co-expression analyses results are only available for *F3H* and *FNS I*, because the expression of the *FNS I*-like genes is too low to be analyzed. F3H shows a strong co-expression with genes of the anthocyanin biosynthesis including *CHS*, *CHI*, *F3’H*, *DFR*, *LDOX*, several glycosyl transferase encoding genes, and a potential anthocyanin transporter encoding gene (S5 File). In contrast, *FNS I* shows a strong association with the flavonol biosynthesis most notable is the co-expression with *FLS1* (S6 File).

**Fig 4.**
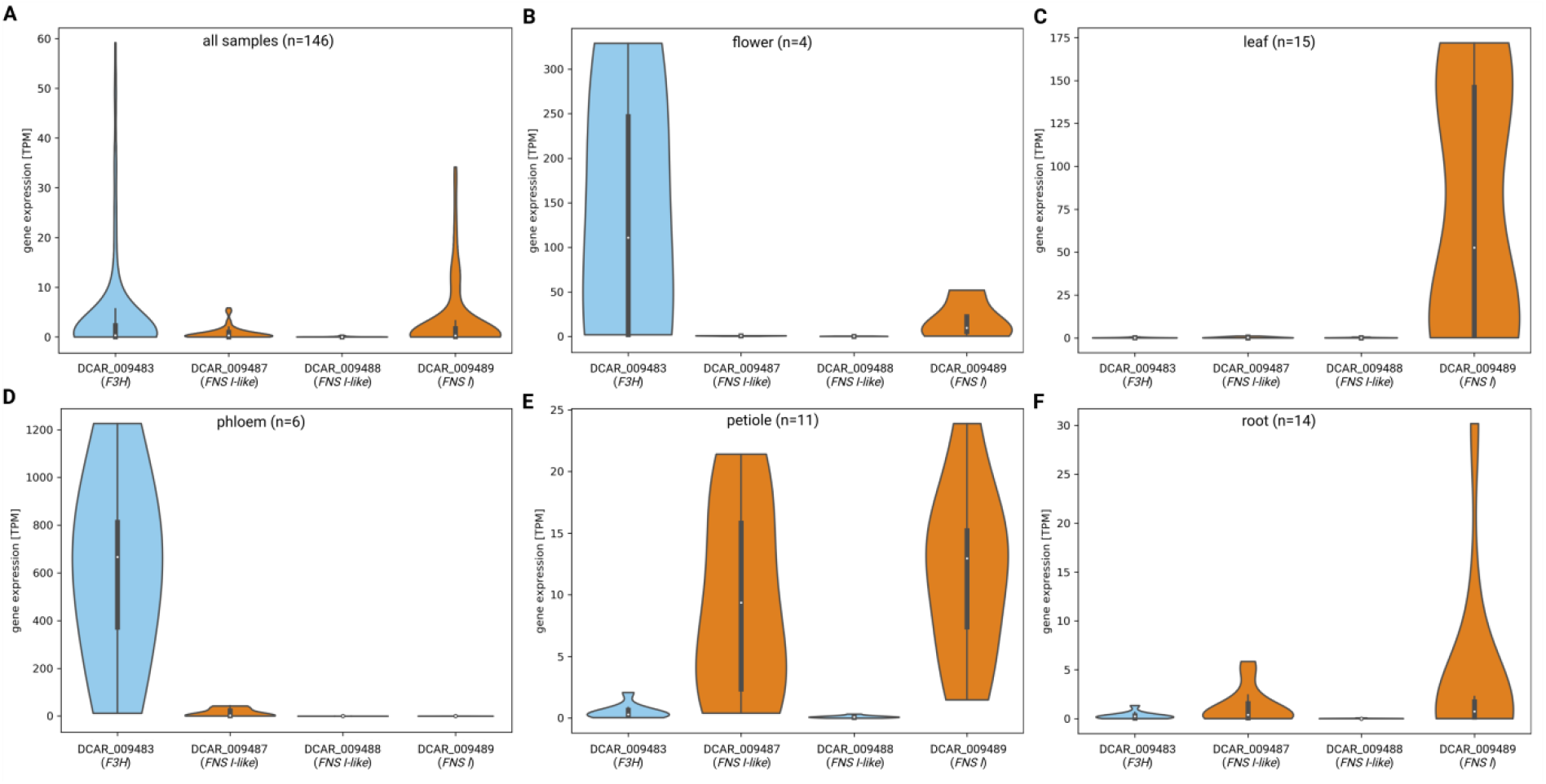
Expression of *F3H*, *FNS I*, and *FNS I*-like genes in carrots. (A) This plot shows the distribution of transcript per million (TPM) values across 146 RNA-seq data sets derived from different tissues/organs and conditions (S2 File). These aggregated expression data reveal that *F3H* (DCAR_009483) and *FNS I* (DCAR_009489) are expressed in several samples, while *FNS I*-like (DCAR_009487) is only weekly expressed in a few selected samples and *FNS I*-like (DCAR_009488) is almost not expressed at all. (B-F) Plots show the expression of the four genes in specific tissues covering flower, leaf, phloem, petiole, and root. This tissue-specific gene expression analysis was restricted to samples with available metadata about the respective sample.

## Discussion

We provide genomic and phylogenetic evidence for the evolution of the Apiaceae *FNS I* from *F3H* through tandem duplication followed by neofunctionalization. These results support a hypothesis about the evolution of *FNS I* from *F3H* [20, 48] and narrow down the initial duplication event. The phylogenetic analysis provides strong support for a single event of Apiaceae FNS I evolution. The nested position of the Apiaceae FNS I clade within the Apiaceae F3H clade is also well supported, while some relationships between F3H sequences of the angiosperms have only low or moderate support.

We show that the F3H duplication most likely took place in a shared ancestor of *D. carota* and *A. graveolens*, and is probably not shared with all members of the Apiacea family as previously hypothesized. Since there is no evidence for this gene duplication in *C. asiatica* which branches early in the Apiaceae, we hypothesize that the F3H duplication took place after the separation of *C. asiatica* from the Daucus/Apium lineage (Fig 3). Additional genome sequences will help to support this hypothesis and to narrow down the precise duplication event within the Apiaceae lineage.

The inspection of conserved amino acid residues in *D. carota* and *A. graveolens* candidate sequences confirmed the presence of one F3H and FNS I in each species. Additionally, both species have at least one sequence that lacks some of the functionally important amino acid residues of a *bona fide* FNS I without having all residues of a F3H (Table 1). This might indicate a different enzymatic activity or promiscuity of these enzymes. It is striking to see that the F3H and FNS I cannot be distinguished based on a P/Y substitution at position 240 (based on *Plagiochasma appendiculatum* PaFNSI/F2H). This difference was previously reported between ancestral FNS I sequences and the sequences of monocot/dicot F3H sequences [8]. All Apiaceae FNS I and F3H sequences show a conserved P at this position suggesting that two independent shifts between F3H and FNS I activity are possible. According to substitution experiments [10], the presence of T106, F131, and E195 in DCAR_009488 indicates that this enzyme has at least some basal FNS activity. It is possible that FNS I-like enzymes have multiple activities [8]. Based on the diagnostic amino acid residues alone, we cannot tell whether (1) these sequences have lost their FNS function in secondary events or (2) represent intermediates in the evolution from F3H towards FNS I. However, the incorporation of additional residues places these sequences in a sister clade to FNS I (Fig 2). This suggests that residues shared between FNS I and FNS I-like sequences were probably present in the shared common ancestor (e.g. F131, I200, R216), while additional FNS I specific residues evolved after separation of both lineages. Based on their phylogenetic relationship, we hypothesize that two *FNS I* copies were present in the common ancestor of *D. carota* and *A. graveolens*. One of these copies was again duplicated in *D. carota* after separation of the lineages leading to *D. carota* and *A. graveolens* hence explaining the presence of three copies in *D. carota*. The preservation of these sequences since the separation of both species indicates a relevance of these FNS I-like sequences. The expression analysis suggests that these genes are active in specific tissues like petiole and root where *FNS I* is also active.

The physical clustering of *FNS I* and *F3H* in the genome could be due to the recent tandem duplication. However, it could be interesting to investigate whether this clustering does also provide an evolutionary benefit. Biosynthetic gene clusters (BGCs) were previously described in numerous plant species [49, 50]. These BGCs are often associated with an evolutionary young trait that provides a particular advantage e.g. in the defense against a pathogen [49]. Given the relevance of flavones in the defense against pathogens [7, 51], it seems possible that the flavone biosynthesis could be a similar trait that evolved in the Apiaceae.

*FNS I* genes were also discovered in a small number of non-Apiaceae species [11, 13, 14]. However, these genes belong to an independent *FNS I* lineage [8]. As more high quality genome sequences of seed plants are released, a systematic search for additional non-Apiaceae *FNS I* sequences could become feasible in the near future. The number of independent FNS I origins remains unknown. Exploration and comparison of additional *FNS I* lineages across plants has the potential to advance our understanding of enzyme evolution.

## Conclusions

In conclusion here we uncovered the duplication mechanism that gave rise to *FNS I* within the Apiaceae family. The gene probably evolved from a tandem duplication of F3H followed by neofunctionalization. The origin of Apiacea *FNS I* appears to be independent from *FNS I* genes described in *Arabidopsis thaliana*, *Oryza sativa*, and *Zea mays*.

## Declarations

### Ethics approval and consent to participate

Not applicable.

### Consent for publication

Not applicable.

### Availability of data and materials

All datasets underlying this study are publicly available or included within the additional files. Scripts developed for this work are freely available on github: https://github.com/bpucker/ApiaceaeFNS1.

### Authors’ contribution

BP and MI performed the analyses and wrote the manuscript.

## Acknowledgements

Many thanks to the German network for bioinformatics infrastructure (de.NBI, grant 031A533A) and the Bioinformatics Resource Facility (BRF) at the Center for Biotechnology (CeBiTec) at Bielefeld University for providing an environment to perform the computational analyses. We used bioRender.com for the construction of some figures. We thank Hanna Marie Schilbert for discussion and comments on the manuscript.

## Supporting information

**S1 File**. Collection of F3H, FNS I, and FNS I-like sequences that were used for the analyses of this study.

**S2 File**. Gene expression values (TPMs) of *Daucus carota F3H*, *FNS I*, and *FNS I-like* genes.

**S3 File**. Phylogenetic trees of F3H, FNS I, and FNS I-like sequences. (A) Constructed by RAxML based on a MAFFT alignment, (B) constructed by RAxML based on a MUSCLE5 alignment, (C) Constructed by FastTree2 based on a MAFFT alignment, (D) Constructed by FastTree2 based on a MUSCLE5 alignment, (E) constructed by IQ-TREE based on a MAFFT alignment, (F) constructed by IQ-TREE based on a MUSCLE5 alignment, (G) constructed by MEGA based on a MAFFT alignment, and (H) constructed by MEGA based on a MUSCLE5 alignment.

**S4 File**. Pairwise comparison of F3H and FNS I sequences. Percentage of identical amino acid residues is displayed in this matrix.

**S5 File**. Results of a co-expression analysis of *F3H* (DCAR_009483) in *Daucus carota* based on 146 RNA-seq data sets.

**S6 File**. Results of a co-expression analysis of *FNS I* (DCAR_009489) in *D. carota* based on 146 RNA-seq data sets.

## References

1. Winkel-Shirley B. Biosynthesis of flavonoids and effects of stress. Current Opinion in Plant Biology. 2002;5:218–23.

2. Nakabayashi R, Mori T, Saito K. Alternation of flavonoid accumulation under drought stress in Arabidopsis thaliana. Plant Signal Behav. 2014;9:e29518.

3. Winkel-Shirley B. Flavonoid Biosynthesis. A Colorful Model for Genetics, Biochemistry, Cell Biology, and Biotechnology. Plant Physiology. 2001;126:485–93.

4. Grotewold E. The genetics and biochemistry of floral pigments. Annu Rev Plant Biol. 2006;57:761–80.

5. Todd JJ, Vodkin LO. Pigmented Soybean (Glycine max) Seed Coats Accumulate Proanthocyanidins during Development. Plant Physiology. 1993;102:663–70.

6. Emiliani J, Grotewold E, Falcone Ferreyra ML, Casati P. Flavonols protect Arabidopsis plants against UV-B deleterious effects. Mol Plant. 2013;6:1376–9.

7. Jiang N, Doseff AI, Grotewold E. Flavones: From Biosynthesis to Health Benefits. Plants. 2016;5:27.

8. Li D-D, Ni R, Wang P-P, Zhang X-S, Wang P-Y, Zhu T-T, et al. Molecular Basis for Chemical Evolution of Flavones to Flavonols and Anthocyanins in Land Plants. Plant Physiol. 2020;184:1731–43.

9. Martens S, Forkmann G. Cloning and expression of flavone synthase II from Gerbera hybrids. The Plant Journal. 1999;20:611–8.

10. Gebhardt YH, Witte S, Steuber H, Matern U, Martens S. Evolution of Flavone Synthase I from Parsley Flavanone 3β-Hydroxylase by Site-Directed Mutagenesis. Plant Physiol. 2007;144:1442–54.

11. Lee YJ, Kim JH, Kim BG, Lim Y, Ahn J-H. Characterization of flavone synthase I from rice. BMB Rep. 2008;41:68–71.

12. Bredebach M, Matern U, Martens S. Three 2-oxoglutarate-dependent dioxygenase activities of Equisetum arvense L. forming flavone and flavonol from (2S)-naringenin. Phytochemistry. 2011;72:557–63.

13. Han X-J, Wu Y-F, Gao S, Yu H-N, Xu R-X, Lou H-X, et al. Functional characterization of a Plagiochasma appendiculatum flavone synthase I showing flavanone 2-hydroxylase activity. FEBS Lett. 2014;588:2307–14.

14. Falcone Ferreyra ML, Emiliani J, Rodriguez EJ, Campos-Bermudez VA, Grotewold E, Casati P. The Identification of Maize and Arabidopsis Type I FLAVONE SYNTHASEs Links Flavones with Hormones and Biotic Interactions. Plant Physiol. 2015;169:1090–107.

15. Wang Q-Z, Downie SR, Chen Z-X. Genome-wide searches and molecular analyses highlight the unique evolutionary path of flavone synthase I (FNSI) in Apiaceae. Genome. 2018;61:103–9.

16. Prescott AG, Stamford NPJ, Wheeler G, Firmin JL. In vitro properties of a recombinant flavonol synthase from Arabidopsis thaliana. Phytochemistry. 2002;60:589–93.

17. Schilbert HM, Schöne M, Baier T, Busche M, Viehöver P, Weisshaar B, et al. Characterization of the Brassica napus Flavonol Synthase Gene Family Reveals Bifunctional Flavonol Synthases. Frontiers in Plant Science. 2021;12.

18. Kawai Y, Ono E, Mizutani M. Evolution and diversity of the 2–oxoglutarate-dependent dioxygenase superfamily in plants. The Plant Journal. 2014;78:328–43.

19. Andersen TB, Hansen NB, Laursen T, Weitzel C, Simonsen HT. Evolution of NADPH-cytochrome P450 oxidoreductases (POR) in Apiales – POR 1 is missing. Molecular Phylogenetics and Evolution. 2016;98:21–8.

20. Martens S, Forkmann G, Britsch L, Wellmann F, Matern U, Lukačin R. Divergent evolution of flavonoid 2-oxoglutarate-dependent dioxygenases in parsley 1. FEBS Letters. 2003;544:93–8.

21. Iorizzo M, Ellison S, Senalik D, Zeng P, Satapoomin P, Huang J, et al. A high-quality carrot genome assembly provides new insights into carotenoid accumulation and asterid genome evolution. Nat Genet. 2016;48:657–66.

22. Jiang Z, Tu L, Yang W, Zhang Y, Hu T, Ma B, et al. The chromosome-level reference genome assembly for Panax notoginseng and insights into ginsenoside biosynthesis. Plant Communications. 2021;2:100113.

23. Li M-Y, Feng K, Hou X-L, Jiang Q, Xu Z-S, Wang G-L, et al. The genome sequence of celery (Apium graveolens L.), an important leaf vegetable crop rich in apigenin in the Apiaceae family. Hortic Res. 2020;7:1–10.

24. Pootakham W, Naktang C, Kongkachana W, Sonthirod C, Yoocha T, Sangsrakru D, et al. De novo chromosome-level assembly of the Centella asiatica genome. Genomics. 2021;113:2221–8.

25. Goodstein DM, Shu S, Howson R, Neupane R, Hayes RD, Fazo J, et al. Phytozome: a comparative platform for green plant genomics. Nucleic Acids Res. 2012;40 Database issue:D1178–86.

26. Pucker B, Reiher F, Schilbert HM. Automatic Identification of Players in the Flavonoid Biosynthesis with Application on the Biomedicinal Plant Croton tiglium. Plants. 2020;9:1103.

27. Downie SR, Katz-Downie DS, Watson MF. A phylogeny of the flowering plant family Apiaceae based on chloroplast DNA rpl16 and rpoC1 intron sequences: towards a suprageneric classification of subfamily Apioideae. Am J Bot. 2000;87:273–92.

28. Stanke M, Keller O, Gunduz I, Hayes A, Waack S, Morgenstern B. AUGUSTUS: ab initio prediction of alternative transcripts. Nucleic Acids Res. 2006;34 suppl_2:W435–9.

29. Pucker B, Holtgräwe D, Weisshaar B. Consideration of non-canonical splice sites improves gene prediction on the Arabidopsis thaliana Niederzenz-1 genome sequence. BMC Research Notes. 2017;10:667.

30. Robinson JT, Thorvaldsdóttir H, Winckler W, Guttman M, Lander ES, Getz G, et al. Integrative Genomics Viewer. Nat Biotechnol. 2011;29:24–6.

31. Gertz EM, Yu Y-K, Agarwala R, Schäffer AA. Altschul SF. Composition-based statistics and translated nucleotide searches: Improving the TBLASTN module of BLAST. BMC Biology. 2006;4:41.

32. Dobin A, Davis CA, Schlesinger F, Drenkow J, Zaleski C, Jha S, et al. STAR: ultrafast universal RNA-seq aligner. Bioinformatics. 2013;29:15–21.

33. Haak M, Vinke S, Keller W, Droste J, Rückert C, Kalinowski J, et al. High Quality de Novo Transcriptome Assembly of Croton tiglium. Front Mol Biosci. 2018;5.

34. Pucker B. Apiaceae FNS I. https://github.com/bpucker/ApiaceaeFNS1. 2022.

35. Price MN, Dehal PS, Arkin AP. FastTree 2 – Approximately Maximum-Likelihood Trees for Large Alignments. PLOS ONE. 2010;5:e9490.

36. Katoh K, Standley DM. MAFFT Multiple Sequence Alignment Software Version 7: Improvements in Performance and Usability. Mol Biol Evol. 2013;30:772–80.

37. Edgar RC. High-accuracy alignment ensembles enable unbiased assessments of sequence homology and phylogeny. 2022;:2021.06.20.449169.

38. Kozlov AM, Darriba D, Flouri T, Morel B, Stamatakis A. RAxML-NG: a fast, scalable and user-friendly tool for maximum likelihood phylogenetic inference. Bioinformatics. 2019;35:4453–5.

39. Kalyaanamoorthy S, Minh BQ, Wong TKF, von Haeseler A, Jermiin LS. ModelFinder: fast model selection for accurate phylogenetic estimates. Nat Methods. 2017;14:587–9.

40. Minh BQ, Schmidt HA, Chernomor O, Schrempf D, Woodhams MD, von Haeseler A, et al. IQ-TREE 2: New Models and Efficient Methods for Phylogenetic Inference in the Genomic Era. Molecular Biology and Evolution. 2020;37:1530–4.

41. Tamura K, Stecher G, Kumar S. MEGA11: Molecular Evolutionary Genetics Analysis Version 11. Mol Biol Evol. 2021;38:3022–7.

42. Tang H, Krishnakumar V, Li J. jcvi: JCVI utility libraries (v0.5.7). 2015. https://doi.org/10.5281/zenodo.31631.

43. Altschul SF, Gish W, Miller W, Myers EW, Lipman DJ. Basic local alignment search tool. J Mol Biol. 1990;215:403–10.

44. Leinonen R, Sugawara H, Shumway M, on behalf of the International Nucleotide Sequence Database Collaboration. The Sequence Read Archive. Nucleic Acids Research. 2011;39 suppl_1:D19–21.

45. Bray NL, Pimentel H, Melsted P, Pachter L. Near-optimal probabilistic RNA-seq quantification. Nat Biotechnol. 2016;34:525–7.

46. Pucker B, Holtgräwe D, Sörensen TR, Stracke R, Viehöver P, Weisshaar B. A De Novo Genome Sequence Assembly of the Arabidopsis thaliana Accession Niederzenz-1 Displays Presence/Absence Variation and Strong Synteny. PLOS ONE. 2016;11:e0164321.

47. Rempel A, Pucker B. BioInfToolServer. BioInfToolServer. 2022. http://pbb.bot.nat.tu-bs.de/KIPEs/. Accessed 31 May 2022.

48. Gebhardt Y, Witte S, Forkmann G, Lukacin R, Matern U, Martens S. Molecular evolution of flavonoid dioxygenases in the family Apiaceae. Phytochemistry. 2005;66:1273–84.

49. Polturak G, Osbourn A. The emerging role of biosynthetic gene clusters in plant defense and plant interactions. PLOS Pathogens. 2021;17:e1009698.

50. Polturak G, Liu Z, Osbourn A. New and emerging concepts in the evolution and function of plant biosynthetic gene clusters. Current Opinion in Green and Sustainable Chemistry. 2022;33:100568.

51. Du Y, Chu H, Wang M, Chu IK, Lo C. Identification of flavone phytoalexins and a pathogen-inducible flavone synthase II gene (SbFNSII) in sorghum. J Exp Bot. 2010;61:983–94.

