## Supplementary figures and images for "Apiaceae *FNS I* originated from *F3H* through tandem gene duplication"

### S3 File

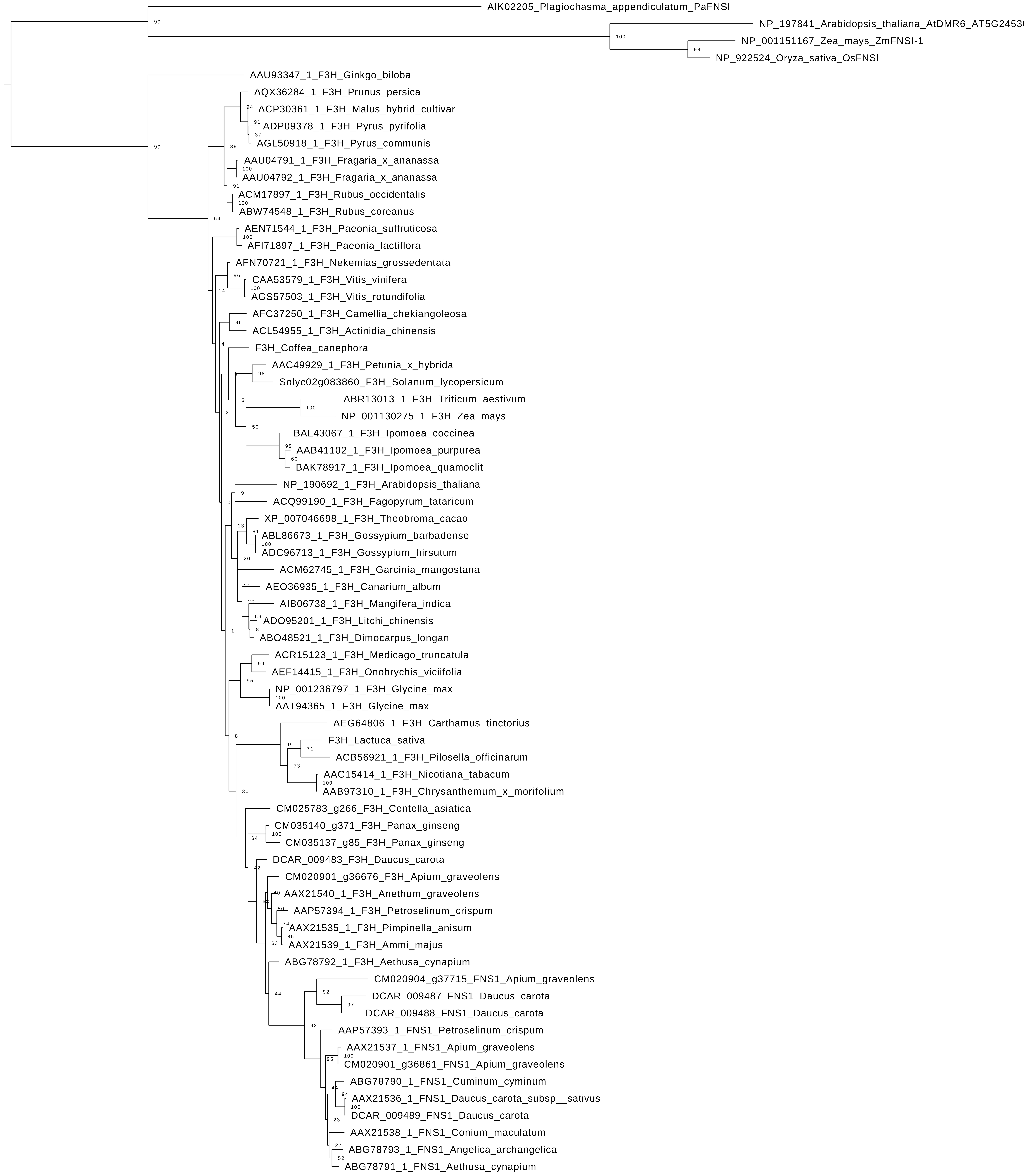

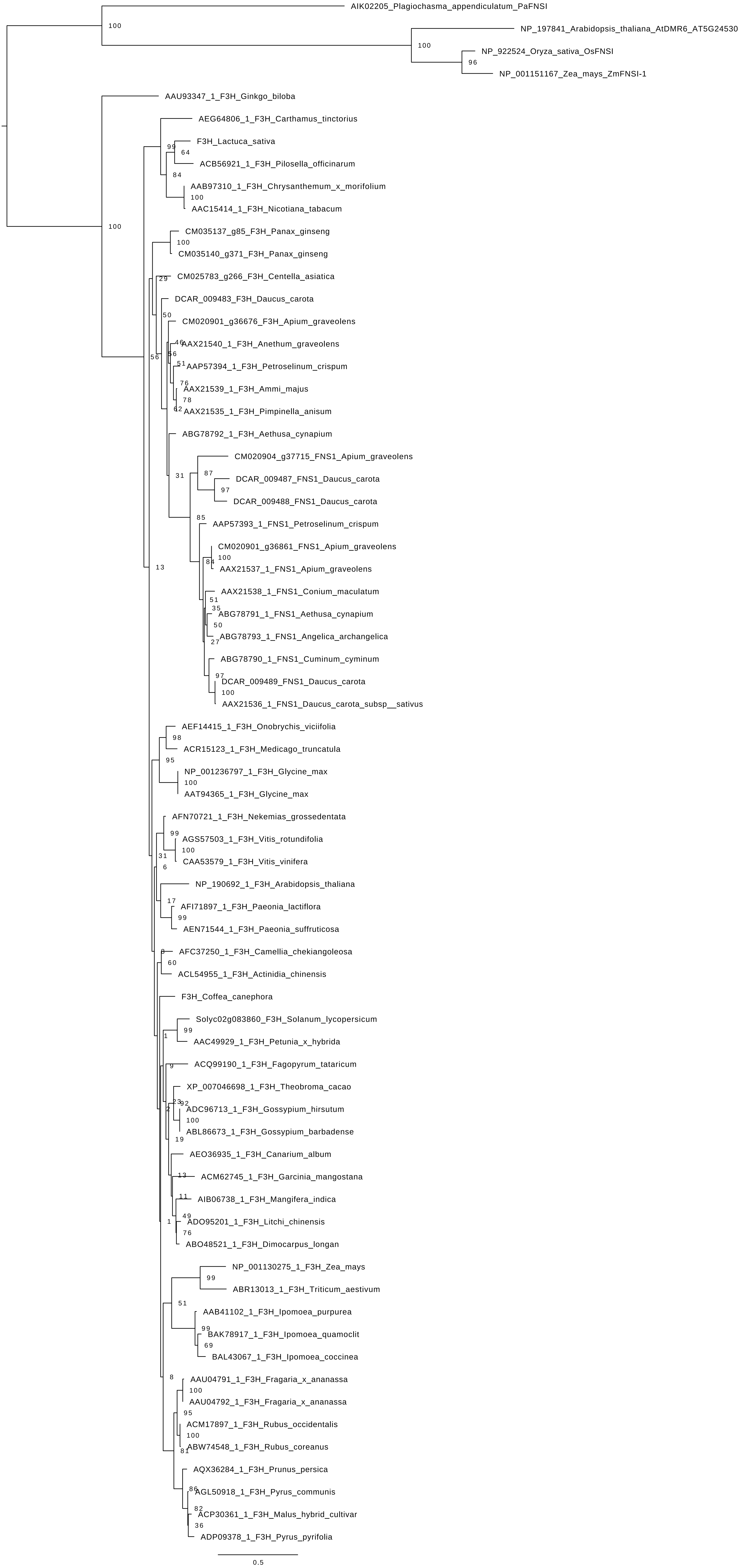

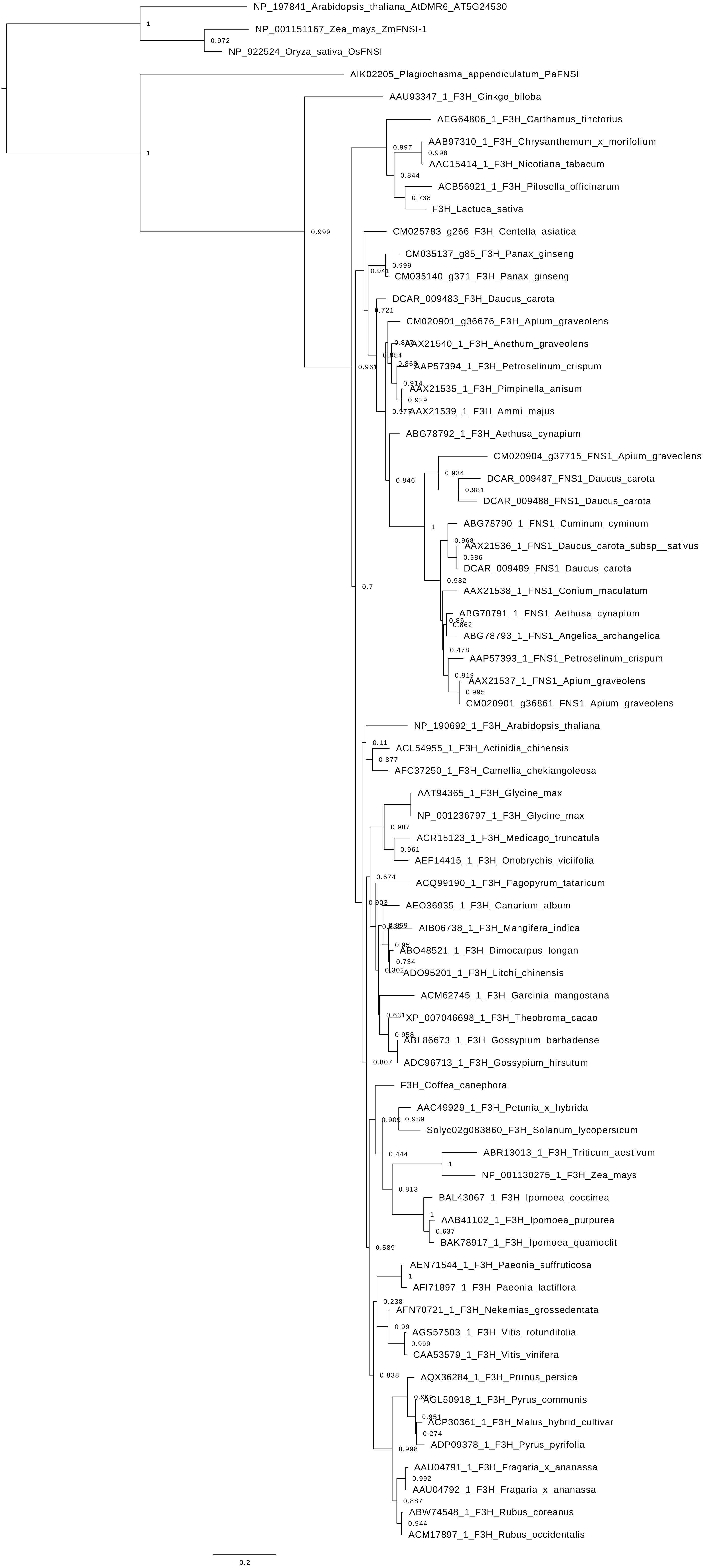

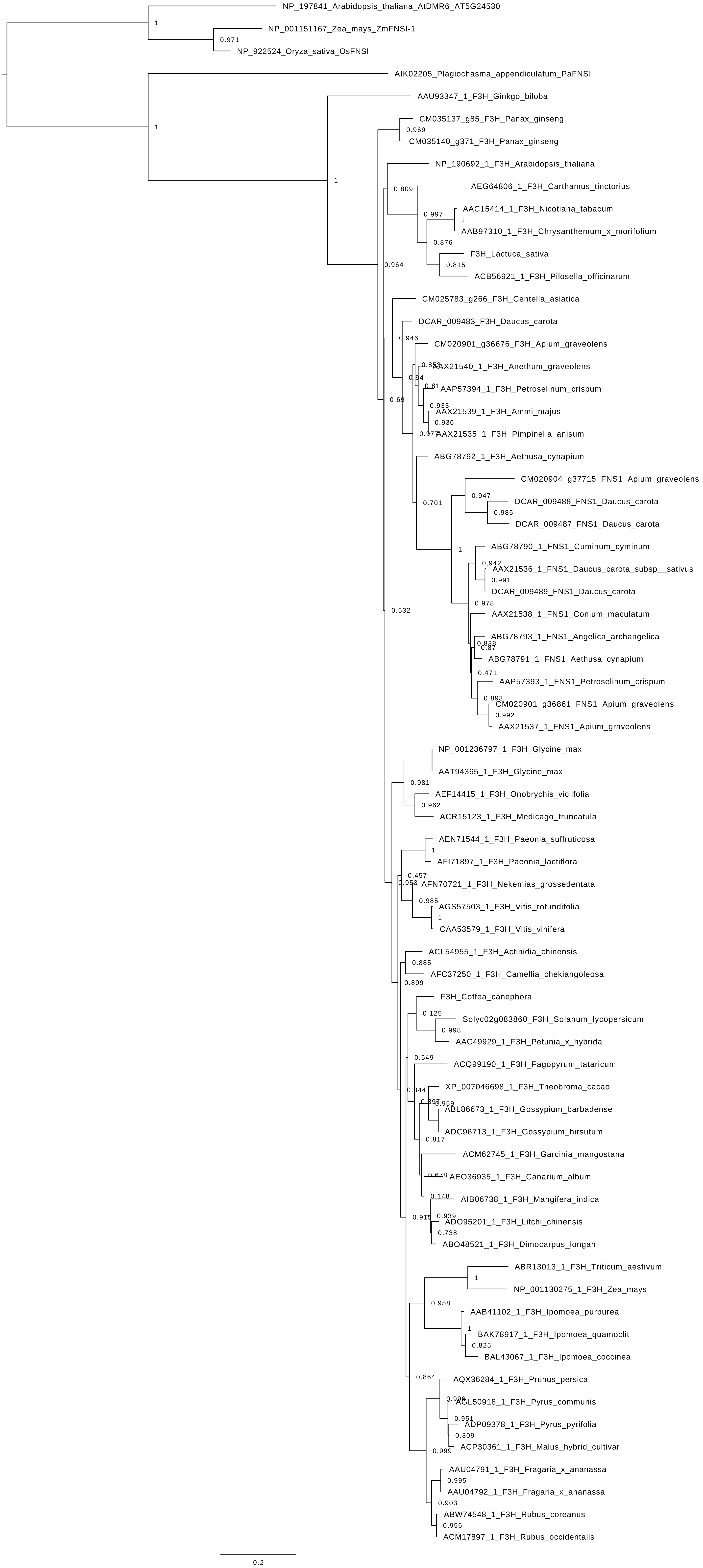

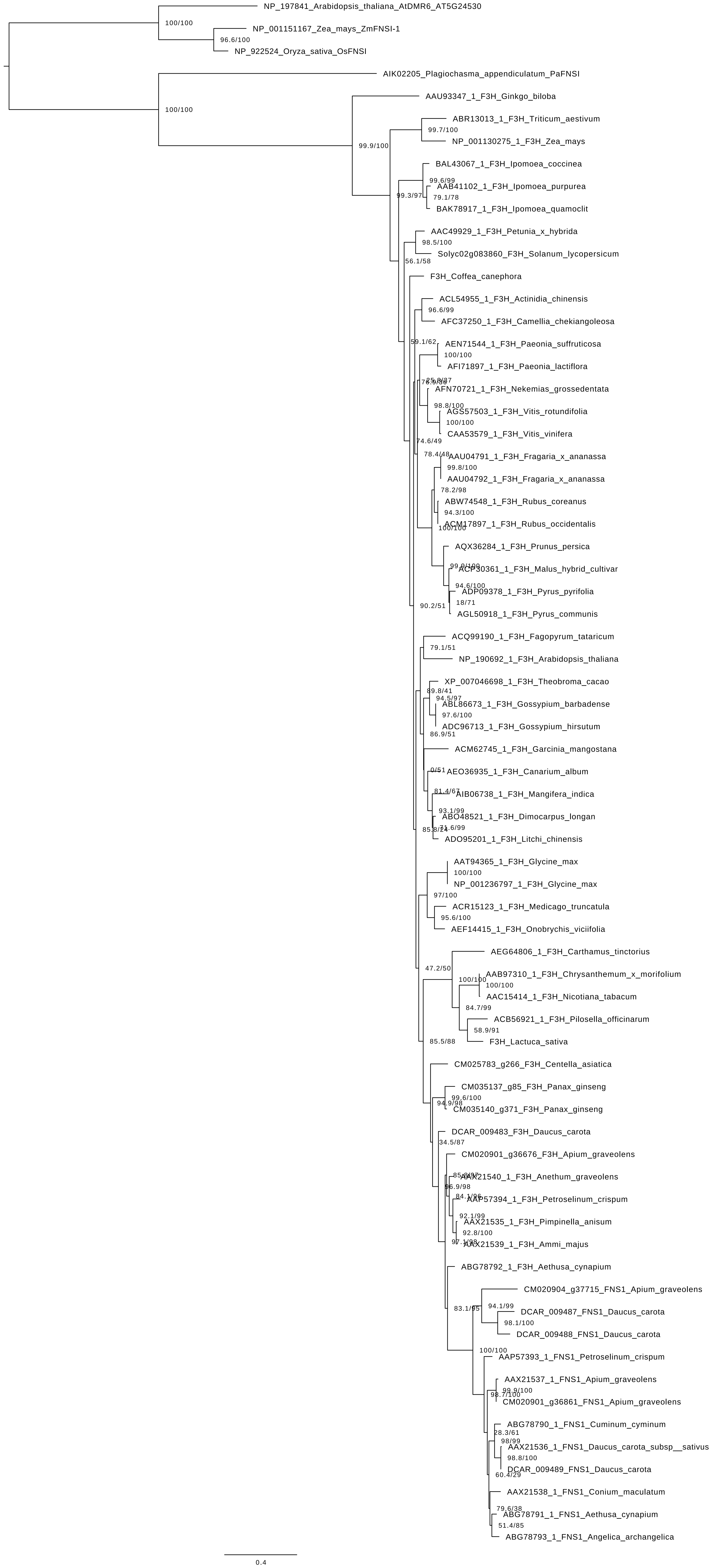

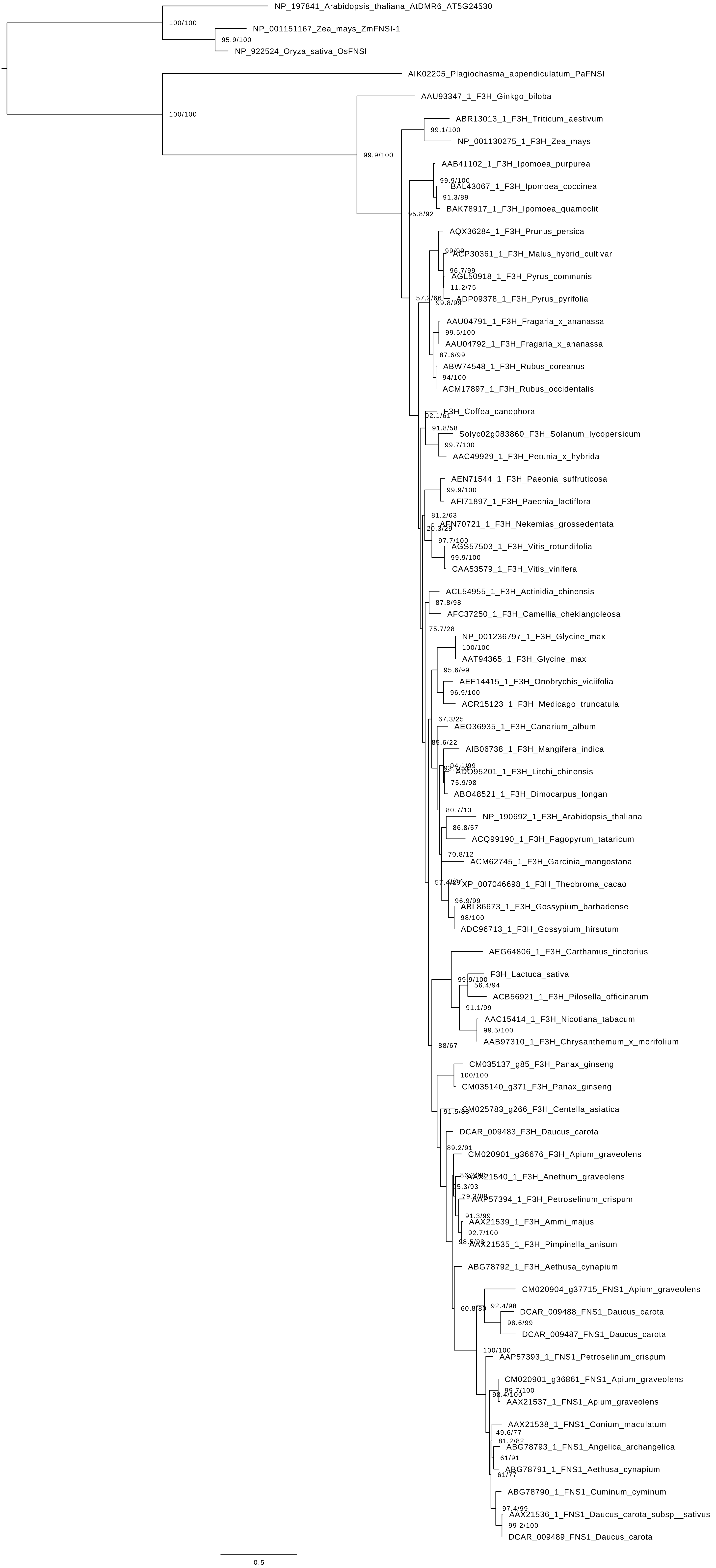

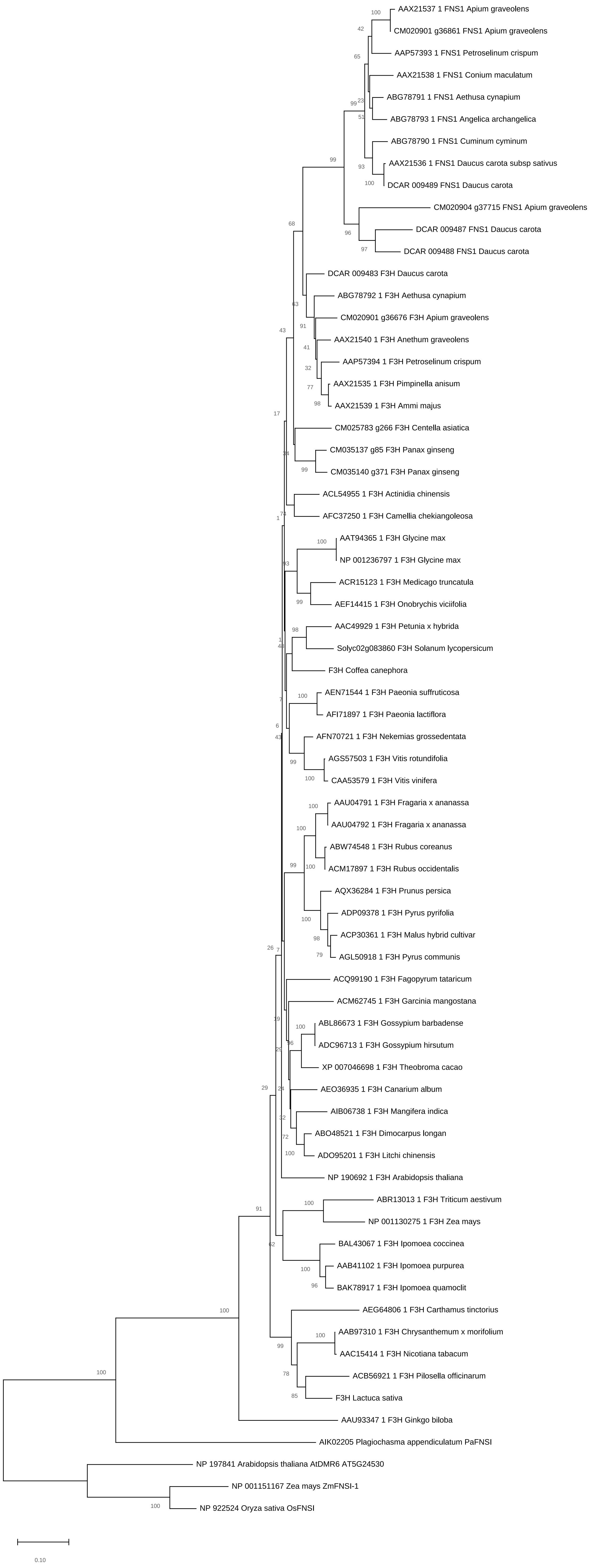

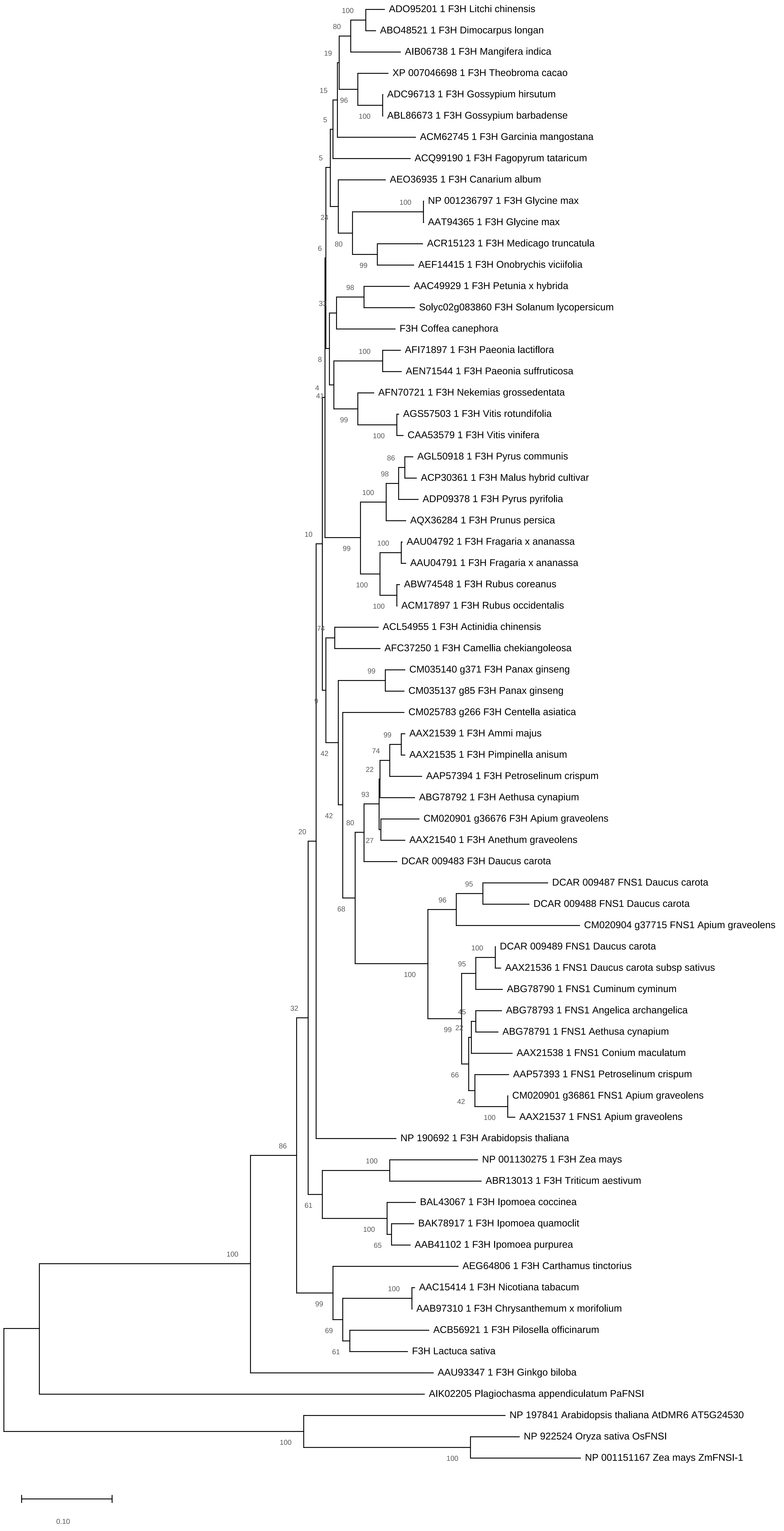
